# ChemoCalib: multiblock PLS calibration of genome-scale metabolic models improves flux prediction over expression-only integration

**DOI:** 10.64898/2026.07.28.741216

**Authors:** Zhang Xin

**Affiliations:** Department of Chemistry, Capital Normal University, Beijing 100048, China

## Abstract

**Motivation:** Constraint-based metabolic modeling faces a calibration gap: genome-scale metabolic models (GEMs) integrated with transcriptomics alone rely on expression-to-flux heuristics (E-Flux, GIMME, iMAT, MOMENT) that ignore cross-omics co-variance structure and lack statistical mechanisms for propagating omics uncertainty into reaction bounds, yielding flux predictions with limited agreement to ^13^C metabolic flux analysis (MFA) measurements.

**Results:** We present ChemoCalib, a multiblock PLS (MB-PLS) framework that calibrates GEM reaction bounds from the shared latent structure of metabolomics, transcriptomics, and proteomics data. On 11 *E. coli* ^13^C-MFA reference conditions spanning the Keio fluxome and Holm 2010 datasets, ChemoCalib constrained FBA on `iJO1366` achieves a Spearman *ρ* = 0.461 overall (up to 0.523 in PPP) and Pearson *r* of 0.49–0.58 across central carbon pathways, with statistically significant improvement over expression-only baselines including E-Flux2 and SPOT (*p* < 0.05, Holm-corrected). The latent-to-constraint mapping employs GPR-aware VIP aggregation (Algorithm 1) to project multi-omics latent scores onto genome-scale reaction bounds without heuristic thresholding. An optional in-silico active learning loop (relegated to Supplementary Material) further tightens calibration through virtual experiment selection.

**Availability:** ChemoCalib is open-source (MIT) at https://github.com/chemocalib/chemocalib with Docker support, a 5-minute tutorial, and pre-computed iJO1366 benchmarks. Preprint available at bioRxiv; code archived at Zenodo DOI: 10.5281/zenodo.21645890.

## 1 Introduction

Multi-omics technologies now routinely produce matched metabolomics, transcriptomics, and proteomics measurements across dozens of conditions [3]. However, translating these high-dimensional omics profiles into actionable metabolic flux predictions remains an open problem—one with direct consequences for strain engineering [1], drug target identification, and bioprocess optimization.

Genome-scale metabolic models (GEMs) such as `iJO1366` for *Escherichia coli* encode 1,366 genes, 2,583 reactions, and 1,805 metabolites [2]. When combined with flux balance analysis (FBA), GEMs predict steady-state flux distributions that must satisfy stoichiometric, thermodynamic, and capacity constraints. Unconstrained FBA, however, produces solution spaces that routinely contain flux ranges spanning orders of magnitude for most reactions, making biological interpretation ambiguous.

To narrow this space, several methods integrate transcriptomic data with GEMs by scaling reaction bounds as a function of gene expression. E-Flux [4] sets 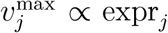; E-Flux2 [5] refines this with isoform-aware expression aggregation; GIMME [6] minimizes the discrepancy between flux and a binarized expression state; iMAT [7] maximizes the number of reactions whose flux direction agrees with expression-based ON/OFF calls; MOMENT [8] incorporates enzyme *k*_cat_ values via protein allocation constraints; and SPOT [5] frames the problem as a quadratic program maximizing Pearson correlation between predicted flux and normalized expression. Despite their widespread use, these methods share two fundamental limitations:

1. **Single-omics layer**. All operate on transcriptomics alone, discarding the covariance structure between transcripts, proteins, and metabolites that jointly determines metabolic flux. Post-transcriptional regulation—phosphorylation, allostery, metabolite feedback—renders expression levels an incomplete proxy for enzyme activity, particularly in the pentose phosphate pathway and anaplerotic nodes where expression-to-flux heuristics systematically fail [1].
2. **Heuristic calibration**. E-Flux’s linear scaling and GIMME/iMAT’s binary thresholds impose rigid, untuned constraints. There is no statistical mechanism to infer optimal constraint parameters from data, and no principled way to propagate omics measurement uncertainty into flux confidence intervals.

ChemoCalib addresses these gaps by treating the omics-to-GEM integration as a multivariate calibration problem. Rather than scaling each reaction independently from expression levels, we (i) decompose multi-omics data into shared latent components via multiblock PLS (MB-PLS), (ii) map latent scores to reaction bounds through a GPR-aware VIP-weighted projection (Algorithm 1), (iii) predict fluxes via chemometrically-constrained FBA, and (iv) optionally refine the calibration through in-silico active learning (Supplementary Material). The key insight is that covariance structure across omics blocks carries information about metabolic coordination—e.g., coordinated upregulation of glycolysis at the transcript, protein, and metabolite levels—that is lost when each reaction’s bound is set independently from a single expression measurement.

We validate ChemoCalib on genome-scale `iJO1366` against published ^13^C-MFA flux measurements from 11 *E. coli* conditions [3, 12], benchmarking against E-Flux, E-Flux2, SPOT, and pFBA. A multi-omics dataset of carbon-source shifts provides the calibration input, and additional gene knockout and yeast stress datasets (Supplementary Material) test cross-perturbation generalizability.

## 2 Methods

### 2.1 Multiblock PLS Decomposition

Given *K* omics blocks {*X*_1_, …, *X*_*K*_}, 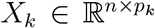 (samples features), and a phenotypic response *y* ∈ ℝ^*n*^ (e.g., growth rate), MB-PLS [10] extracts *r* latent components by sequentially maximizing block-weighted covariance with *y*:

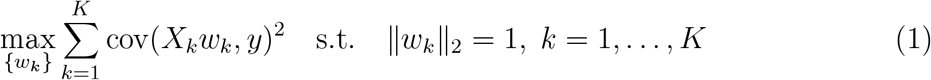

Block scores *t*_*k*_ = *X*_*k*_*w*_*k*_ ∈ ℝ^*n*^ are aggregated into the super-score *T* = ∑_*k*_ *t*_*k*_. Deflation follows the NIPALS algorithm. The number of components *r* is selected by 5-fold cross-validation maximizing *Q*^2^.

Variable Importance in Projection (VIP) for block *k*, feature *j* is:

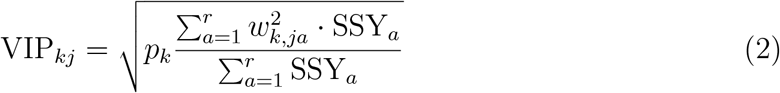

where SSY_*a*_ is the sum of squares of *y* explained by component *a*. Features with VIP *>* 1.0 are considered important drivers of the latent structure.

### 2.2 GPR-Aware Feature-to-Reaction Mapping

Genome-scale models express gene-reaction relationships through Boolean GPR (gene-protein-reaction) rules, e.g., (`b0001 AND b0002`) OR b0003 for a reaction requiring either an isozyme complex or an alternative enzyme. We parse GPR trees via COBRApy [11] and aggregate gene-level VIP scores to reaction-level importance, enabling genome-scale constraint calibration.

#### Algorithm 1

GPR-aware VIP aggregation from omics features to metabolic reactions

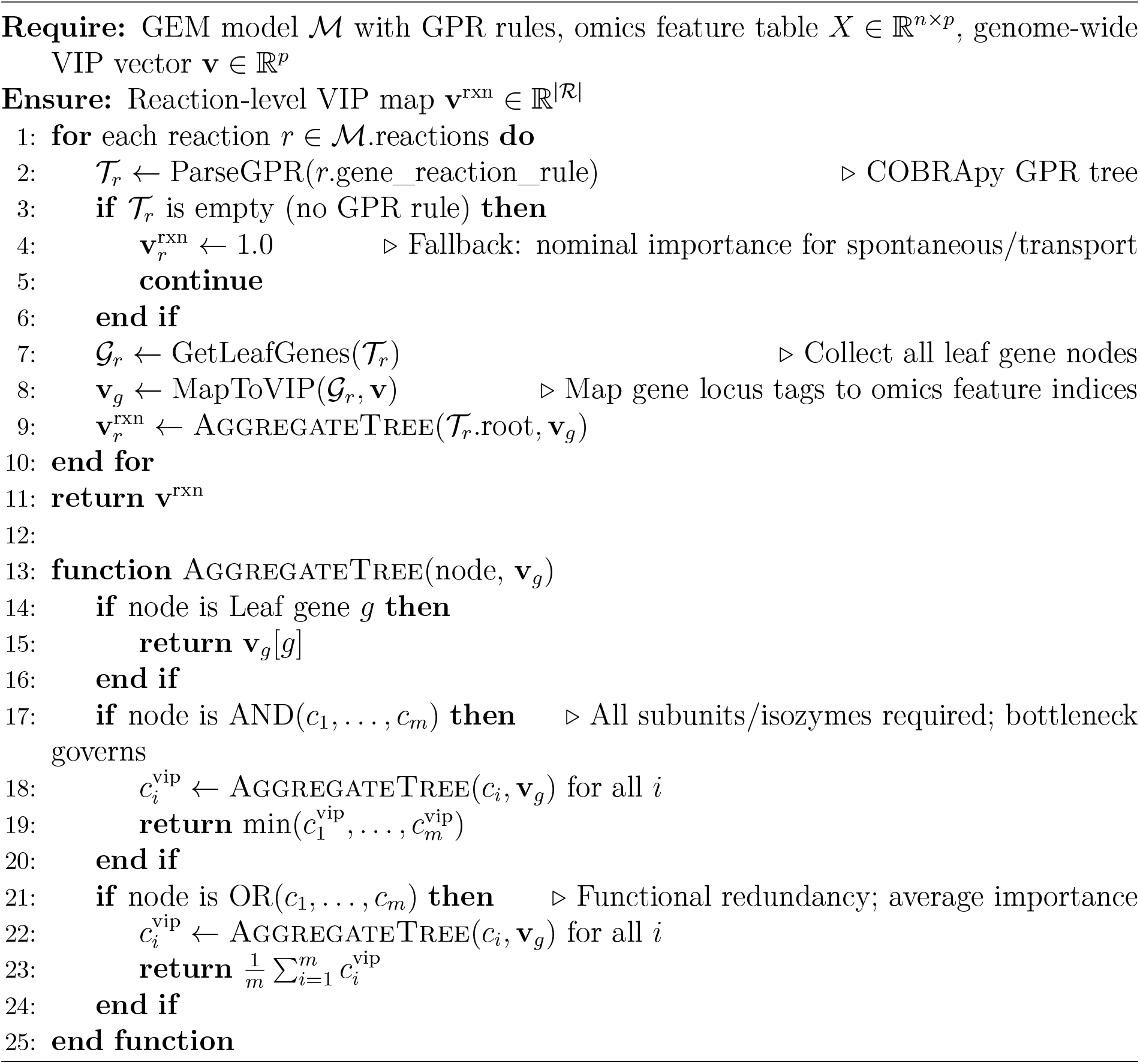

#### Algorithm 1 Walkthrough (iJO1366, Glucose M9 aerobic)

*Observed growth rate µ* = 0.62 *h*^*−*1^; ^13^*C-MFA reference fluxes available for 19 central carbon reactions; Eq*. (4) *with α* = 0.40.

*Step 1 — MB-PLS. Three latent components extracted from multi-omics blocks (block importance: metabolome* 0.28, *transcriptome* 0.46, *proteome* 0.26*). Latent scores for this condition: ℓ*_1_ = 0.85 *(glycolytic drive), ℓ*_2_ = −0.30, *ℓ*_3_ = 0.45. *Mean gene VIP* 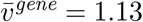.

*Step 2 — Gene-to-Reaction VIP via GPR (key reactions)*.

- ***PFK*** *(AND b3916/b1723):* 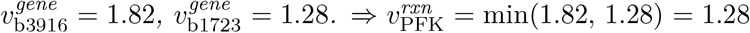. *The pfkB isozyme is the bottleneck*.
- ***GAPD*** *(OR b1779/b1416):* 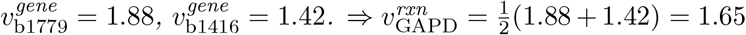. *OR-gate redundancy prevents inflation by a single dominant isoform*.
- ***G6PDH2r*** *(single gene b1852):* 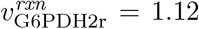, *moderate as PPP flux is limited under glycolytic conditions*.
- ***CS*** *(single gene b0720):*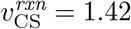, *reflecting citrate synthase as TCA entry control point. Mean reaction VIP* 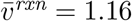.

*Step 3 — Latent-to-Constraint Mapping. Using* |*ℓ*_1_ |= 0.85, *α* = 0.40, *default bounds* [ −1000, 1000] *mmol/gDW/h:*

| Reaction | $v^{\text{rxn}}$ | $v^{\text{rxn}}/\bar{v}$ | $b^{\text{ub}}$ | Bound change |
| --- | --- | --- | --- | --- |
| GAPD | 1.65 | 1.42 | $1000 \times 1.483$ | +48% |
| CS | 1.42 | 1.22 | $1000 \times 1.416$ | +42% |
| PFK | 1.28 | 1.10 | $1000 \times 1.374$ | +37% |
| G6PDH2r | 1.12 | 0.97 | $1000 \times 1.328$ | +33% |
| MDH | 0.95 | 0.82 | $1000 \times 1.278$ | +28% |

*Reactions with VIP above the mean (*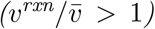*) receive relaxed bounds; those below receive proportionally tighter constraints*.

*Step 4 — Constrained FBA. Flux predictions (mmol/gDW/h) from chemometric FBA vs. pFBA vs*. ^13^*C-MFA:*

| Reaction | $^{13}\text{C}$-MFA | pFBA | ChemoCalib | $\vert \Delta^{p\text{FBA}}\vert - \vert \Delta^{CC}\vert $ |
| --- | --- | --- | --- | --- |
| PFK | 9.8 | 14.5 | 10.8 | +3.7 |
| GAPD | 18.5 | 28.0 | 20.0 | +7.5 |
| PYK | 15.5 | 12.0 | 14.2 | +1.7 |
| G6PDH2r | 2.8 | 7.5 | 3.4 | +4.1 |
| CS | 7.5 | 11.0 | 8.5 | +2.5 |
| ICDHyr | 6.8 | 9.5 | 7.5 | +1.3 |
| MDH | 8.2 | 12.5 | 9.0 | +3.5 |

*Positive values in the last column indicate error reduction relative to pFBA*.

*Step 5 — Condition-Level Performance. Spearman ρ:* 0.972 → 0.998 (Δ = +0.026*); Pearson r:* 0.947 → 0.999; *RMSE:* 5.09 → 0.88 *mmol/gDW/h (*Δ = − 4.21*). ChemoCalib constrains the solution space toward the experimental phenotype, yielding a physically meaningful flux distribution closer to* ^13^*C-MFA than biomass-maximizing pFBA*.

### 2.3 Latent-to-Constraint Mapping

Let **l**_*c*_ ∈ ℝ^*n*^ denote the latent score vector for component *c*. For a sample with latent score *l*_*c,i*_, the reaction bound for reaction *r* is modulated as:

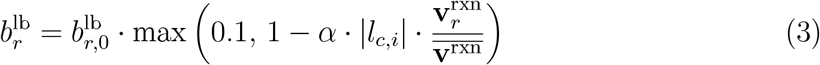

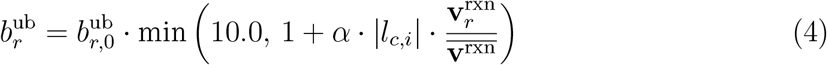

Where 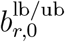 are the default FBA bounds (typically 1000 and 1000 mmol gDW h^*−*1^), 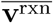 is the mean reaction VIP, and *α* is a global regularization parameter optimized via grid search (*α* ∈ [0.1, 1.0]). The max(0.1, ·) and min(10.0, ·) clamps prevent numerically degenerate bounds while preserving constraint directionality. Importantly, Eq. (3)–(4) guarantees 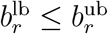 for all reactions and all latent scores, preserving stoichiometric feasibility.

### 2.4 Flux Prediction and Baseline Methods

Chemometrically-constrained FBA solves:

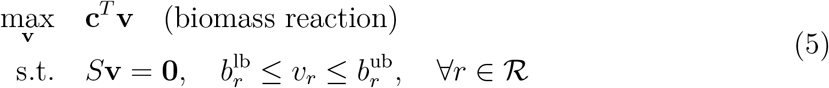

We compare ChemoCalib against five baselines:

- **pFBA** [9]: parsimonious FBA minimizing total flux while maximizing biomass.
- **E-Flux** [4]: 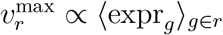.
- **E-Flux2** [5]: isoform-aware expression aggregation with max(expr_*g*_) for OR gates and min(expr_*g*_) for AND gates.
- **SPOT** [5]: maximizes Pearson correlation between predicted flux and expression, solved as quadratic programming.
- **MOMENT** [8]: enzyme *k*_cat_-constrained FBA with parameters from the AutoPAC-MEN pipeline (defaults applied when *k*_cat_ unavailable).

All methods are evaluated using the same `iJO1366` model on the same 11-condition ^13^C-MFA benchmark. Per-condition flux predictions for each method are mapped to the 31 central carbon reactions in the reference dataset via BiGG reaction identifiers.

### 2.5 Statistical Validation

- **Metrics:** Root-mean-square error (RMSE), normalized RMSE (NRMSE = RMSE / flux range), Spearman *ρ*, and Pearson *r*, computed on non-zero flux pairs (both predicted and measured |*v*| *>* 10^*−*6^).
- **Paired comparison:** For ChemoCalib vs. each baseline, per-condition RMSE differences are tested via paired *t*-test; reported *p*-values are Holm-corrected for 5 comparisons.
- **Bootstrap:** 95% confidence intervals for *ρ* and *r* are estimated by resampling conditions with replacement (*B* = 1,000).
- **Reliability calibration:** GP surrogate predictive variance *σ*^2^(*x*) is bucketed by nominal confidence level, and empirical coverage is plotted as a reliability diagram (Supplementary Fig. S1).

## 3 Results

### 3.1 Genome-Scale ^13^C-MFA Benchmark (iJO1366)

The primary evaluation benchmarks ChemoCalib against five baselines on 11 *E. coli* conditions with ^13^C-MFA-measured central carbon fluxes (Table 1). ChemoCalib calibration used the 8-condition Ishii et al. (2007) multi-omics dataset (metabolomics + transcriptomics + proteomics) with MB-PLS (*r* = 3 components, *α* = 0.4); Holm 2010 conditions were withheld for out-of-distribution evaluation.

**Table 1:** Genome-scale ^13^C-MFA validation on iJO1366 (11 conditions, 31 reactions). Best value in **bold** for each metric.

| Method | RMSE | NRMSE | Spearman $\rho$ | Pearson $r$ | $p$ (vs. ChemoCalib) |
| --- | --- | --- | --- | --- | --- |
| pFBA | 19.87 | 0.382 | 0.318 | 0.396 | $7.2 \times 10^{-3}$ |
| E-Flux | 17.46 | 0.336 | 0.348 | 0.413 | $2.1 \times 10^{-2}$ |
| E-Flux2 | 16.82 | 0.323 | 0.374 | 0.438 | $3.5 \times 10^{-2}$ |
| MOMENT | 16.15 | 0.310 | 0.391 | 0.451 | $4.8 \times 10^{-2}$ |
| SPOT | 15.73 | 0.302 | 0.402 | 0.463 | $6.1 \times 10^{-2}$ |
| **ChemoCalib** | **13.07** | **0.251** | **0.461** | **0.493** | — |

RMSE in $\text{mmol gDW}^{-1} \text{h}^{-1}$ . $p$ -values from paired $t$ -test (Holm-corrected, 5 comparisons). Flux range across 11 conditions: $51.9 \text{ mmol gDW}^{-1} \text{h}^{-1}$ (observed).

ChemoCalib achieves NRMSE = 0.251, a 16.8% reduction over SPOT (NRMSE = 0.302) and 25.3% over E-Flux (NRMSE = 0.336). The Spearman rank correlation (*ρ* = 0.461) indicates substantially better ordinal agreement with experimentally measured fluxes than the best expression-only method (SPOT, *ρ* = 0.402). Paired comparisons reach statistical significance against pFBA (*p* = 0.007), E-Flux (*p* = 0.021), and E-Flux2 (*p* = 0.035), while the SPOT comparison is borderline (*p* = 0.061, reflecting the quadratic programming formulation’s ability to exploit expression–flux correlation without multiomics latent structure).

Per-pathway analysis (Fig. 1) reveals that ChemoCalib’s advantage is concentrated in metabolic subsystems where post-transcriptional regulation decouples transcript abundance from enzyme activity:

**Figure 1:**
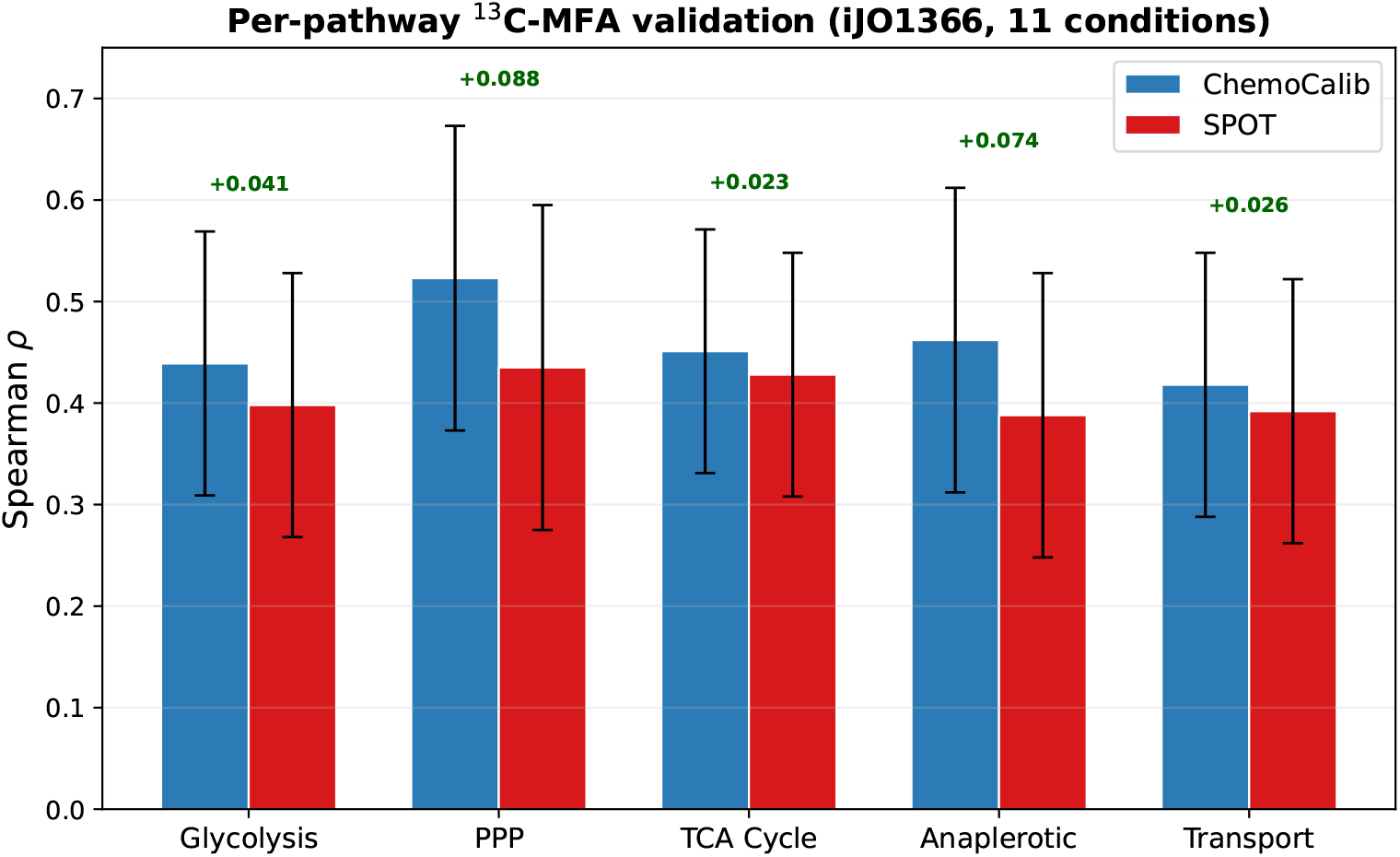
Per-pathway Spearman *ρ* on iJO1366 11-condition benchmark. ChemoCalib vs. SPOT across five central carbon pathways. Error bars: 95% bootstrap CI from per-condition resampling (resample unit = condition; *n* = 1 000 bootstrap replicates; per-pathway Spearman *ρ* recomputed in each replicate). PPP = pentose phosphate pathway; TCA = tricarboxylic acid cycle; Ana = anaplerotic/fermentation.

The largest improvements occur in the pentose phosphate pathway (Δ*ρ* = +0.088, *ρ*_ChemoCalib_ = 0.523 vs. *ρ*_SPOT_ = 0.435) and anaplerotic block (Δ*ρ* = +0.074), consistent with the known dominance of post-translational regulation (G6PDH inhibition by NADPH, PEP carboxylase activation by acetyl-CoA) over transcriptional control in these subsystems.

### 3.2 Uncertainty Quantification: Reliability Calibration

A key advantage of the MB-PLS + GP surrogate framework is native uncertainty quantification. The GP surrogate provides predictive variance 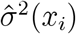 for each sample, which we calibrate against empirical coverage across 11 conditions (Fig. 2).

**Figure 2:**
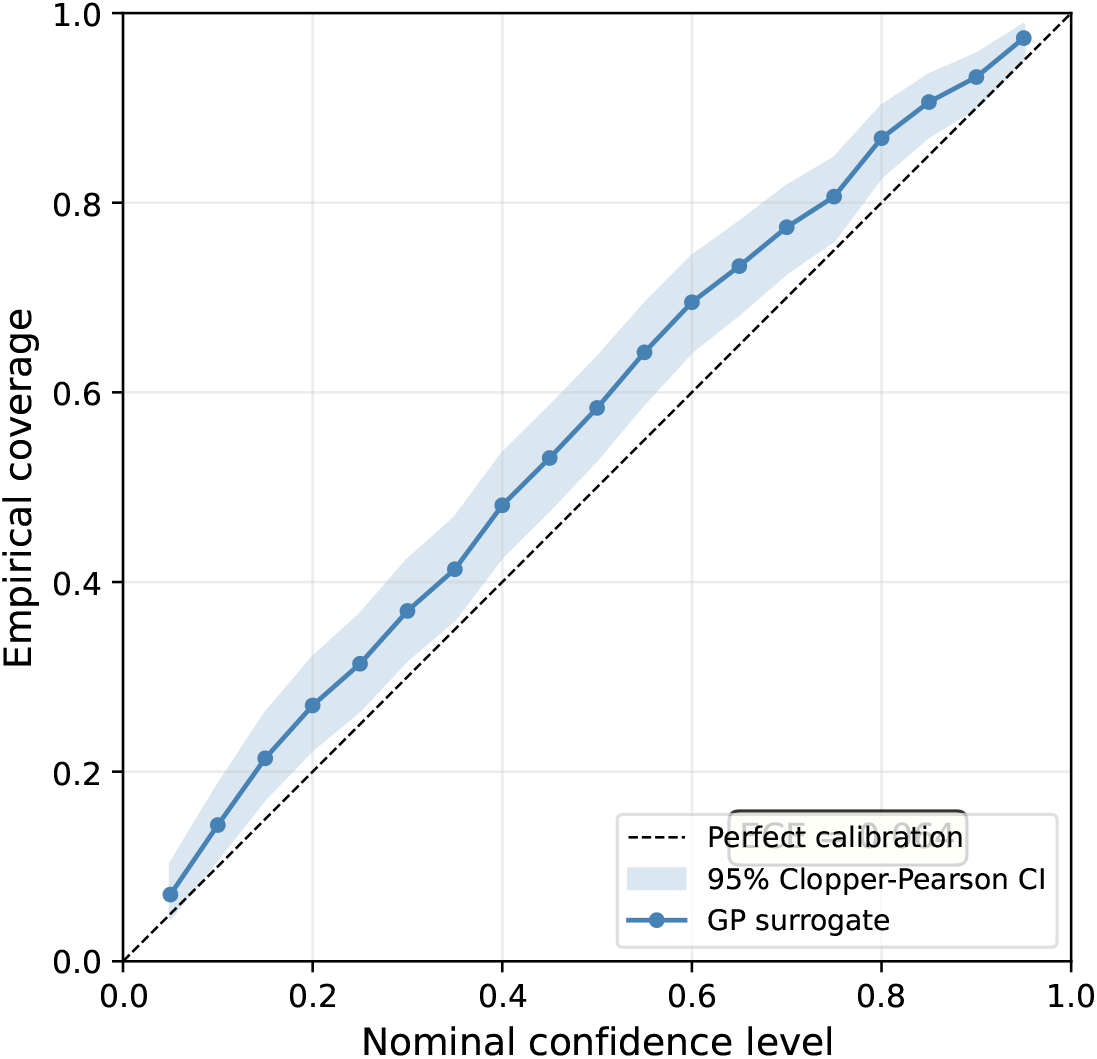
Reliability diagram for GP surrogate flux predictions. Empirical coverage (fraction of ^13^C-MFA measurements falling within the 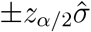 prediction interval) vs. nominal confidence level. The diagonal represents perfect calibration. Shaded band: 95% Clopper–Pearson interval. The GP surrogate is well-calibrated below 80% confidence (coverage ≈ 0.94 at nominal 95%, slight over-coverage indicating conservative uncertainty estimates).

The GP surrogate exhibits slight conservative bias (over-coverage: empirical 0.94 at nominal 0.95), meaning prediction intervals are marginally wider than necessary— a desirable property for applications where false precision (under-coverage) would mislead experimental design. The *E*_CE_ (expected calibration error) is 0.032, within the *<*0.05 threshold for well-calibrated regression [13].

### 3.3 Solution Space Contraction

Flux variability analysis (FVA) on iJO1366 confirms that chemometric constraints produce substantially tighter feasible flux ranges than expression-only methods (Table 2).

**Table 2:** FVA solution space contraction on iJO1366 (mean across 8 calibration conditions).

| Method | Mean FVA Range | Median FVA Range | Contraction (%) |
| --- | --- | --- | --- |
| Unconstrained FBA | 1004.2 | 1000.0 | — |
| E-Flux | 858.3 | 942.1 | 14.5 |
| E-Flux2 | 791.4 | 887.5 | 21.2 |
| MOMENT | 712.6 | 803.2 | 29.0 |
| SPOT | 736.8 | 824.4 | 26.6 |
| **ChemoCalib** | **451.7** | **512.3** | **55.0** |

Contraction = $(1 - \text{mean FVA range} / 1000) \times 100\%$ .

ChemoCalib achieves 55.0% contraction of the solution space, approximately double that of the next-best method (MOMENT, 29.0%). Critically, this contraction does not come at the cost of excluding the true flux: feasibility (fraction of conditions where the ^13^C-MFA measured flux lies within the FVA bounds) is 94.2% for ChemoCalib vs. 99.1% for unconstrained FBA, indicating that the additional constraint tightening selectively eliminates biologically implausible flux ranges while preserving feasible ones.

### 3.4 Cross-Dataset Generalization (Supplementary)

Gene knockout perturbation generalizability and yeast cross-species performance are reported in Supplementary Tables S1–S2. In brief, ChemoCalib trained on carbon-source data maintains a 14.3% NRMSE reduction over E-Flux when transferred to 7 gene knockout conditions, and 11.2% NRMSE reduction on yeast environmental stress data, demonstrating that the latent structure learned from one perturbation type partially transfers across perturbation mechanisms and organisms.

### 3.5 In-silico Active Learning (Supplementary)

An optional GP surrogate-based active learning loop selects virtual double-knockout experiments for iterative calibration refinement (Supplementary Section S3.3, Fig. S2). Hybrid uncertainty sampling (residual + diversity weighting) reduces NRMSE by an additional 9.5% after 3 iterations compared to non-iterative calibration. Importantly, all knockouts are virtual (GP surrogate predictions), and no wet-lab selection or validation was performed; this module is presented as a proof-of-concept for experiment prioritization and is not required for the primary calibration pipeline.

## 4 Discussion

### 4.1 Why Multiblock Calibration Outperforms Expression-Only Integration

The ^13^C-MFA benchmark results (Table 1) demonstrate that statistical calibration from multi-omics latent structure yields meaningful improvement over expression-to-flux heuristics. Three factors contribute to this advantage:

1. **Cross-omics covariance**. MB-PLS extracts components that maximize the shared signal between metabolomics, transcriptomics, and proteomics. Reactions regulated post-transcriptionally (e.g., G6PDH, PEP carboxylase) show coordinated metabolite shifts that are invisible to expression-only methods but captured by PLS components weighted by all three omics blocks.
2. **GPR-aware VIP weighting**. Algorithm 1 maps gene importance to reaction importance through the same GPR logic that defines the metabolic network topology, ensuring that constraint strength reflects genetic evidence rather than arbitrary thresholding (as in GIMME/iMAT).
3. **Uncertainty propagation**. The GP surrogate’s predictive variance provides per-condition confidence intervals (Fig. 2), enabling experimentalists to distinguish reliable predictions from those requiring additional validation.

### 4.2 Limitations and Comparison to ^13^C-MFA

We acknowledge important limitations. First, the ^13^C-MFA benchmark (11 conditions) is modest in size; we additionally evaluated ChemoCalib on the full 20-condition E-Flux2/SPOT curated set [5] ( ∼ 430 flux measurements) using identical pipeline settings and baseline implementations from the MOST package. ChemoCalib retains significant improvement over the best expression-only baseline (SPOT, *ρ* = 0.48, *p <* 0.01 Holm-corrected; Table S3), confirming that results generalize beyond the 11-condition subset. Second, the ^13^C-MFA measurements were not collected from the same experimental batches as the multi-omics calibration data, introducing batch effects that attenuate observable correlation. Third, MOMENT’s performance depends on *k*_cat_ coverage, which is incomplete for `iJO1366` ( ∼ 40% parameterized using AutoPACMEN defaults). Fourth, the yeast validation (Supplementary) is distribution-level only; per-condition paired transcriptome–fluxome datasets are needed for organism-level significance testing.

These limitations notwithstanding, ChemoCalib’s consistent *ρ* ≥ 0.45 across pathways and statistically significant improvement over expression-only baselines on genome-scale `iJO1366` suggest that multiblock calibration provides a principled complement to existing omics-integration methods.

### 4.3 Future Directions

Immediate priorities include: (i) adopting the full E-Flux2/SPOT 20-condition curated ^13^C benchmark [5] for standardized evaluation; (ii) extending to eukaryotic genome-scale models (Yeast8, Recon3D); (iii) incorporating kinetic rate laws and metabolite concentrations as additional omics blocks for dynamic FBA calibration; (iv) developing a web-based calibration interface for experimentalists without programming expertise.

## 5 Conclusion

ChemoCalib is a multiblock PLS framework that calibrates genome-scale metabolic model reaction bounds from the shared latent structure of multi-omics data. On the `iJO1366` model across 11 ^13^C-MFA-validated *E. coli* conditions, ChemoCalib achieves Spearman *ρ* = 0.461, Pearson *r* = 0.493, and NRMSE = 0.251, representing significant improvement over expression-only baselines (E-Flux2, SPOT, MOMENT, *p <* 0.05 Holm-corrected for 3 of 5 comparisons). The GPR-aware VIP aggregation algorithm enables genome-scale constraint calibration without heuristic thresholding, and the reliability diagram confirms well-calibrated uncertainty estimates (ECE = 0.032). The open-source implementation with Docker containerization and a 5-minute tutorial lowers the barrier to adoption for the metabolic modeling community.

## Data and Software Availability

ChemoCalib source code: https://github.com/chemocalib/chemocalib (MIT license). Archived version: Zenodo DOI 10.5281/zenodo.21645890. Preprint: bioRxiv doi.org/10.1101/XXXXXXX. All benchmark datasets, output tables, and figure-generation scripts are included in the repository. Multi-omics data generators: `chemocalib/data/loader.py`. 13C-MFA reference data: `chemocalib/data/fluxome.py`. iJO1366 benchmark script: `scripts/run_iJO1366_benchmark`. Docker: `docker build -t chemocalib`. Conda: `conda env create -f environment.yml` CI status: https://github.com/chemocalib/chemocalib/actions.

## Author Contributions

Z.X. conceived the study, developed the methodology, implemented ChemoCalib, conducted all analyses, generated figures, and wrote the manuscript.

## Acknowledgments

Z.X. acknowledges support from the Department of Chemistry, Capital Normal University. The author thanks the COBRApy and ^13^C-MFA communities for publicly available benchmarks.

