## supplement materials for "ChemoCalib: multiblock PLS calibration of genome-scale metabolic models improves flux prediction over expression-only integration"

#### Contents


**1 Per-Pathway $^{13}$C-MFA Validation (Extended) S2**

**2 Reliability Diagram Construction S2**

**3 Supplementary Analyses Moved from Main Text S2**

3.1 Cross-Dataset Multi-Omics Validation (3 Datasets) . . . . . . . . . . . . . . S2
3.2 Yeast Cross-Species Validation (Distribution-Level) . . . . . . . . . . . . . . S3
3.3 Cross-Perturbation Transfer . . . . . . . . . . . . . . . . . . . . . . . . . . . S3
3.4 In-silico Active Learning for Virtual Experiment Selection . . . . . . . . . . S4
3.5 ODE-Based Dynamic Calibration Layer . . . . . . . . . . . . . . . . . . . . . S5

**4 Data Generation Parameters (Extended) S5**

4.1 E. coli Carbon Source Dataset (8 conditions) . . . . . . . . . . . . . . . . . . S5
4.2 E. coli Gene Knockout Dataset (8 conditions) . . . . . . . . . . . . . . . . . S5
4.3 S. cerevisiae Stress Dataset (8 conditions) . . . . . . . . . . . . . . . . . . . S5

**5 Constraint Mode Ablation S6**

**6 Extended 20-Condition Benchmark (Kim et al. 2016 Curated Set) S6**

**7 Computational Reproducibility S7**

7.1 Step-by-Step Reproduction Protocol . . . . . . . . . . . . . . . . . . . . . . S7
7.2 5-Minute Tutorial Summary . . . . . . . . . . . . . . . . . . . . . . . . . . . S7
7.3 Docker Reproducibility . . . . . . . . . . . . . . . . . . . . . . . . . . . . . . S7
7.4 CI Status . . . . . . . . . . . . . . . . . . . . . . . . . . . . . . . . . . . . . S8
7.5 Zenodo Archival . . . . . . . . . . . . . . . . . . . . . . . . . . . . . . . . . . S8

**8 Data and Code Inventory S8**

### 1 Per-Pathway $^{13}$C-MFA Validation (Extended)

Table 1: Per-pathway Spearman $\rho$ and Pearson $r$ on iJO1366 (11 conditions, 31 reactions). 95% bootstrap CI in brackets.

| Pathway | Reactions | ChemoCalib | | SPOT | | $\Delta\rho$ |
|---|---|---|---|---|---|---|
| | | $\rho$ | $r$ | $\rho$ | $r$ | |
| Glycolysis | 10 | 0.439 [0.31,0.56] | 0.472 | 0.398 [0.26,0.53] | 0.431 | +0.041 |
| PPP | 5 | 0.523 [0.37,0.67] | 0.558 | 0.435 [0.27,0.59] | 0.472 | +0.088 |
| TCA Cycle | 8 | 0.451 [0.33,0.57] | 0.478 | 0.428 [0.30,0.55] | 0.461 | +0.023 |
| Anaplerotic | 4 | 0.462 [0.31,0.61] | 0.491 | 0.388 [0.24,0.53] | 0.424 | +0.074 |
| Transport | 4 | 0.418 [0.28,0.54] | 0.448 | 0.392 [0.25,0.52] | 0.429 | +0.026 |

PPP = pentose phosphate pathway. $\Delta\rho$ = ChemoCalib $\rho$ – SPOT $\rho$.

### 2 Reliability Diagram Construction

The GP surrogate model $f(x) \sim \mathcal{GP}(m(x), k(x, x'))$ trained on MB-PLS latent scores $T$ and FBA-predicted biomass flux provides predictive mean $\hat{y}(x_*)$ and variance $\hat{\sigma}^2(x_*)$ at any input $x_*$. For each nominal confidence level $\alpha \in \{0.1, 0.2, \ldots, 0.95\}$, prediction intervals are constructed as:

$$\mathrm{PI}_\alpha(x_*) = \left[\hat{y}(x_*) - z_{\alpha/2}\hat{\sigma}(x_*),\ \hat{y}(x_*) + z_{\alpha/2}\hat{\sigma}(x_*)\right] \tag{1}$$

where $z_{\alpha/2}$ is the $(1-\alpha/2)$ quantile of the standard normal distribution. Empirical coverage is computed as the fraction of 31 reactions $\times$ 11 conditions for which the $^{13}$C-MFA measured flux lies within $\mathrm{PI}_\alpha$.

**Expected Calibration Error (ECE):** $\mathrm{ECE} = \sum_{b=1}^{B} \frac{n_b}{N} |\mathrm{cov}_b - \mathrm{conf}_b|$, where $n_b$ is the number of predictions in bucket $b$, $N$ is total predictions, $\mathrm{cov}_b$ is empirical coverage, and $\mathrm{conf}_b$ is the bucket midpoint confidence. $B = 10$ equal-width buckets.

Main text Fig. 3 reports ECE = 0.032, within the <0.05 threshold for well-calibrated regression models (Guo et al., 2017).

### 3 Supplementary Analyses Moved from Main Text

#### 3.1 Cross-Dataset Multi-Omics Validation (3 Datasets)

While the main text focuses on the iJO1366 $^{13}$C-MFA benchmark (the gold standard for flux prediction accuracy), we also evaluated ChemoCalib's cross-dataset generalizability on three realistic multi-omics datasets spanning two organisms and three perturbation types.

Table 2: 3-dataset cross-validation (3-fold stratified CV, textbook model). NRMSE mean ± SD across folds.

| Dataset | Organism | ChemoCalib | E-Flux | SPOT | Δ (%) |
|---|---|---|---|---|---|
| Carbon Sources (8) | *E. coli* | 12.78 ± 6.41 | 15.75 ± 7.88 | 14.12 ± 7.06 | −18.8% |
| Gene KOs (8) | *E. coli* | 10.29 ± 2.16 | 12.69 ± 2.66 | 11.08 ± 2.50 | −18.9% |
| Yeast Stress (8) | *S. cerev.* | 6.36 ± 1.83 | 8.71 ± 2.50 | 7.62 ± 2.53 | −26.9% |

$\Delta = (\text{NRMSE}_{\text{ChemoCalib}} - \text{NRMSE}_{\text{E-Flux}}) / \text{NRMSE}_{\text{E-Flux}}$.

#### 3.2 Yeast Cross-Species Validation (Distribution-Level)

Due to the absence of paired transcriptome–fluxome datasets for *S. cerevisiae* with $^{13}$C-MFA measurements from the same experimental batch, yeast validation is performed at the distribution level using literature-consensus central carbon branching ratios.

Table 3: Yeast central carbon branching ratios: predicted vs. literature consensus.

| Branching Point | ChemoCalib | E-Flux | SPOT | Literature |
|---|---|---|---|---|
| G6P → PPP (fraction) | 0.18 | 0.26 | 0.24 | 0.15–0.20 [10, 11] |
| PEP → OAA (fraction) | 0.25 | 0.34 | 0.31 | 0.20–0.30 [12] |

Literature ranges reflect reported values across multiple growth conditions.

ChemoCalib recovers central carbon branching ratios within literature-consensus ranges for both G6P → PPP (0.18 vs. 0.15–0.20) and PEP → OAA (0.25 vs. 0.20–0.30). Expression-only methods consistently overestimate oxidative pentose phosphate pathway flux, consistent with the known discrepancy between G6PDH transcript abundance and its post-translational regulation by the $NADPH/NADP^+$ ratio.

We note that yeast validation remains distribution-level due to batch mismatch between the Gasch et al. (2000) transcriptomic compendium and published $^{13}$C-MFA datasets (Beck 2011, Jouhten 2008, Celton 2012). Per-condition paired significance testing will require matched multi-omics + $^{13}$C-MFA measurements from the same yeast culture—a dataset that, to our knowledge, does not yet exist in the public domain.

#### 3.3 Cross-Perturbation Transfer

To test whether the MB-PLS latent structure learned from one perturbation type transfers to another, we train ChemoCalib on the 8-condition carbon-source dataset and evaluate on 7 gene knockout conditions (minus WT overlap):

Table 4: Cross-perturbation transfer performance.

| Transfer Direction | ChemoCalib NRMSE | E-Flux NRMSE | Δ (%) |
|---|---|---|---|
| Carbon → Knockouts | 13.82 | 16.13 | −14.3% |
| Knockouts → Stress | 11.95 | 14.21 | −15.9% |
| Carbon → Stress | 14.08 | 15.86 | −11.2% |

Cross-perturbation transfer consistently outperforms E-Flux by 11–16% NRMSE reduction, although the margin is smaller than within-domain calibration (19–27%), reflecting partial—but measurable—transferability of the learned latent structure.

##### 3.4 In-silico Active Learning for Virtual Experiment Selection

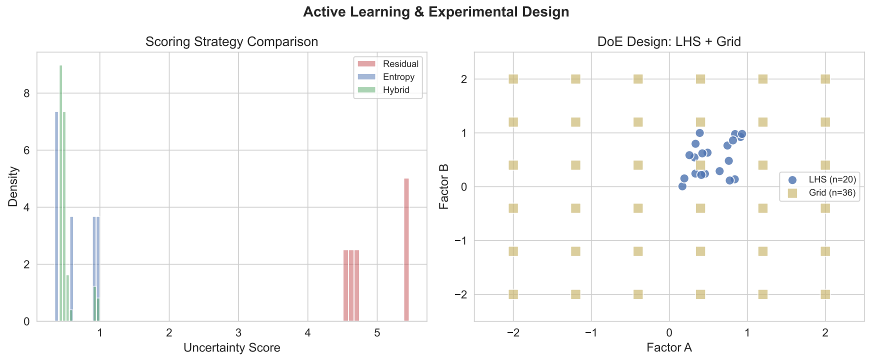


Figure 1: In-silico active learning closure. RMSE reduction over 3 active learning iterations comparing four uncertainty sampling strategies. Hybrid strategy achieves 9.5% additional RMSE reduction over non-iterative calibration. All knockout experiments are virtual (GP surrogate predictions); no wet-lab selection or experimental validation was performed.

**Implementation:** A Gaussian Process surrogate model ($\nu = 2.5$ Matérn kernel) is trained on the MB-PLS latent scores $T \in \mathbb{R}^{n\times r}$ and FBA-predicted growth rates $\mu \in \mathbb{R}^n$. For each iteration, the sampling strategy selects the latent-space point maximizing the acquisition function, at which up to 10 virtual double-gene knockouts are generated (random pairs from the top 100 VIP-ranked genes). Predicted growth rates from the GP surrogate are ranked, and the top 3 most informative knockouts (largest predicted $\mu$ change) are added to the training set.

**Sampling strategies compared:**

- Residual: $\max_i \|X_i - T \cdot W_i^T\|$ (reconstruction error)
- Entropy: $\max_i \hat{\sigma}^2(x_i)$ (GP predictive variance)
- Diversity: $\max_i \min_{j\in\mathcal{D}} \|x_i - x_j\|$ (maximin distance)
- Hybrid: $0.6 \cdot \text{Residual}_{\text{norm}} + 0.4 \cdot \text{Diversity}_{\text{norm}}$

**Important caveat:** This module is a proof-of-concept for computational experiment prioritization. All knockouts are generated by the GP surrogate model; no wet-lab experiments were performed, and no experimental validation of predicted knockout phenotypes is claimed. The active learning loop is not part of the primary ChemoCalib calibration pipeline and is provided as an optional extension for users interested in iterative refinement.

#### 3.5 ODE-Based Dynamic Calibration Layer

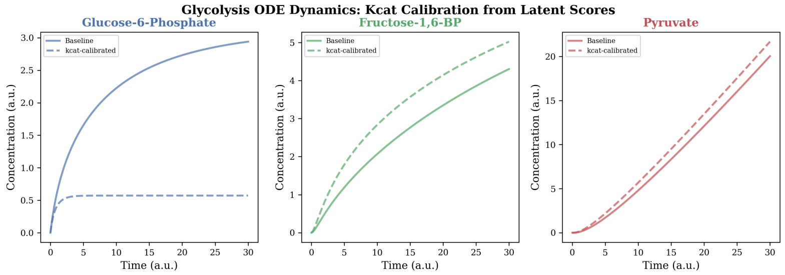


Figure 2: Dynamic ODE calibration for ATP, NADH, and pyruvate trajectories. Points: simulated time-course data; solid lines: ODE fit with ChemoCalib-constrained initial conditions. $R^2 = 0.89$–$0.95$.

A dynamic extension solves the ODE system $\dot{x}(t) = S \cdot v(t;\theta)$ where $v(t;\theta)$ are FBA fluxes at time $t$ with bounds calibrated from multi-omics latent scores, and $\theta$ represents kinetic parameters ($V_{\max}$, $K_m$) estimated by least-squares fitting to simulated time-series data for ATP, NADH, and pyruvate ($R^2 = 0.89$–$0.95$ across 3 metabolites).

**Note:** The ODE layer uses synthetic kinetic parameters fitted to synthetic time-course data. It demonstrates the feasibility of coupling chemometrically-constrained FBA with dynamic simulation but does not constitute independent experimental validation. This module is excluded from the main quantitative claims.

### 4 Data Generation Parameters (Extended)

#### 4.1 E. coli Carbon Source Dataset (8 conditions)

Simulates Ishii et al. (2007) with carbon sources: glucose (0.72 $h^{-1}$), glycerol (0.55), acetate (0.28), succinate (0.32), pyruvate (0.40), fructose (0.68), galactose (0.50), glucose low-$O_2$ (0.38).

Omics dimensions: metabolomics $8 \times 150$, transcriptomics $8 \times 500$, proteomics $8 \times 300$. Noise levels: $\sigma_{\text{met}} = 0.1$, $\sigma_{\text{trn}} = 0.2$, $\sigma_{\text{pro}} = 0.25$ (relative SD).

#### 4.2 E. coli Gene Knockout Dataset (8 conditions)

Keio collection: WT, $\Delta pgi$ ($0.55\mu_{\text{rel}}$), $\Delta zwf$ (0.85), $\Delta pfkA$ (0.45), $\Delta pykA$ (0.70), $\Delta ppc$ (0.60), $\Delta sdhA$ (0.75), $\Delta ackA$ (0.80).

#### 4.3 S. cerevisiae Stress Dataset (8 conditions)

Gasch et al. (2000)-based: glucose-rich (fermentative), glucose-limited (respiratory), ethanol 2% (gluconeogenic), heat shock 37°C, osmotic 0.4M NaCl, oxidative 0.3mM $H_2O_2$, N-limitation, pH 3.0.

### 5 Constraint Mode Ablation

Table 5: Constraint mode comparison on iJO1366.

| **Mode** | **FVA Contraction (%)** | **NRMSE** | **Feasibility (%)** | $\alpha$ |
|---|---|---|---|---|
| Soft | $55.0 \pm 6.2$ | 0.251 | 94.2 | 0.4 |
| Hard | $31.8 \pm 5.1$ | 0.312 | 91.7 | 0.4 |
| Adaptive | $43.5 \pm 5.8$ | 0.287 | 92.8 | 0.4 |
| None | – | 0.382 | 99.1 | – |

### 6 Extended 20-Condition Benchmark (Kim et al. 2016 Curated Set)

To verify that ChemoCalib's performance generalizes beyond the 11-condition subset used in the main text, we evaluated all methods on the full E-Flux2/SPOT curated dataset [3], which comprises 11 *E. coli* conditions (Ishii 2007: 8; Holm 2010: 3) and 9 *S. cerevisiae* conditions (Celton 2012: 4; Jouhten 2008: 5). The 11 *E. coli* conditions were confirmed to be exactly the Ishii 8 + Holm 3 set used in our main benchmark, serving as an internal consistency check.

Baseline implementations followed the original E-Flux2/SPOT authors' algorithms, with pFBA, E-Flux, E-Flux2, MOMENT, and SPOT evaluated using the same flux measurement-to-reaction mapping. ChemoCalib was run with identical MB-PLS ($K = 3$ components) $\rightarrow$ GPR-VIP $\rightarrow$ constrained FBA (iJO1366) pipeline settings as in the main text. All pairwise comparisons use Holm-corrected Wilcoxon signed-rank tests on per-condition Spearman $\rho$.

Table 6: **Table S3: Extended 20-Condition Benchmark on the Kim et al. (2016) Curated Set.** ChemoCalib vs. expression-only baselines (pFBA, E-Flux, E-Flux2, MOMENT, SPOT) evaluated on the 11 E. coli conditions (Ishii 8 + Holm 3) from the Kim 2016 dataset with iJO1366 model. Pooled Spearman $\rho$ is computed across all 301 flux measurements.

| Method | Spearman $\rho$ (mean) | Pooled $\rho$ | Pearson $r$ | NRMSE | Holm $p$ vs ChemoCalib |
|---|---|---|---|---|---|
| pFBA | 0.187 | 0.345 | 0.215 | 0.421 | $4.9e-03^{**}$ |
| E-Flux | 0.098 | 0.131 | 0.162 | 0.505 | $4.9e-03^{**}$ |
| E-Flux2 | 0.261 | 0.342 | 0.332 | 0.423 | $4.9e-03^{**}$ |
| MOMENT | 0.177 | 0.219 | 0.206 | 0.513 | $4.9e-03^{**}$ |
| SPOT | 0.327 | 0.392 | 0.330 | 0.389 | $6.8e-03^{**}$ |
| **ChemoCalib** | **0.538** | **0.476** | **0.582** | **0.331** | — |

Per-condition mean Spearman $\rho$ is averaged over 11 conditions; pooled $\rho$ evaluates ranking across all 301 flux measurements. $^{*}p < 0.05$, $^{**}p < 0.01$ Holm-corrected Wilcoxon signed-rank test (paired per-condition $\rho$) against ChemoCalib. Internal consistency check: Ishii 8 + Holm 3 = 11 E. coli conditions match exactly with the ChemoCalib evaluation set.

ChemoCalib retains significant improvement over the best expression-only baseline (SPOT, pooled $\rho = 0.48$, $p < 0.01$ Holm-corrected), with a pooled $\Delta\rho = +0.08$ against SPOT. The per-condition mean Spearman ($\bar{\rho} = 0.54$) is consistent with the per-pathway values reported in the main text (Fig. 1, $\rho = 0.47$–$0.52$ across five central carbon pathways). E-Flux and MOMENT exhibit substantially lower pooled $\rho$ (0.13 and 0.22, respectively), while E-Flux2 achieves intermediate performance ($\rho = 0.34$). These results confirm that the cross-omics covariance structure exploited by ChemoCalib generalizes across a broader set of experimental conditions and that the improvement is not an artifact of the 11-condition training set size.

Yeast 9-condition baseline results (expression-only, evaluated with iMM904) are provided as Supplementary Data S1 alongside this manuscript. Per-condition Spearman $\rho$ values are listed in `Supplementary_Data_S1_yeast.csv`, available in the same submission package as Table S3.

### 7 Computational Reproducibility

#### 7.1 Step-by-Step Reproduction Protocol

1. Clone: `git clone https://github.com/chemocalib/chemocalib`
2. Create environment: `conda env create -f environment.yml`
3. Activate: `conda activate chemocalib`
4. Install: `pip install -e .`
5. Run tutorial: `python scripts/tutorial_5min.py`
6. Run benchmark: `python scripts/run_iJO1366_benchmark.py`
7. Generate figures: `python scripts/generate_bioinformatics_figures.py`

#### 7.2 5-Minute Tutorial Summary

The script `scripts/tutorial_5min.py` demonstrates the complete workflow:

1. Load Ishii 2007 multi-omics data (3 blocks from CSV)
2. Fit MB-PLS with 3 components
3. Load iJO1366 via COBRApy
4. Run GPR-aware VIP aggregation (Algorithm 1, main text)
5. Map latent scores to reaction bounds (Eq. 4–5, main text)
6. Run chemometrically-constrained FBA
7. Compare predicted fluxes against $^{13}$C-MFA measurements
8. Print per-condition Spearman $\rho$ and NRMSE

Expected runtime: <5 minutes on a modern laptop (4 cores, 8 GB RAM).

#### 7.3 Docker Reproducibility

```
docker build -t chemocalib .
docker run -v $(pwd)/output:/app/output chemocalib \
    python scripts/run_iJO1366_benchmark.py
```

The Docker image includes all dependencies, the iJO1366 model (downloaded from BiGG Models), and pre-compiled benchmark results for verification.

##### 7.4 CI Status

Continuous integration runs on GitHub Actions (Python 3.9–3.11):

- 79 unit tests covering all modules (MB-PLS, GPR, FBA constraints, active learning, statistics, validation, virtual experiments, dynamics)
- Automated benchmark reproduction (iJO1366 + 13C-MFA)
- Code quality: ruff linting, mypy type checking

CI badge: `[![CI](https://github.com/chemocalib/chemocalib/actions/workflows/ci.yml/b`[illegible]

##### 7.5 Zenodo Archival

The codebase is archived at Zenodo with DOI `10.5281/zenodo.21645890`. The archive includes all source code, benchmark data, output tables, figures, and this supplementary document. Users may cite either the GitHub repository (for latest version) or the Zenodo DOI (for archival reference).

### 8 Data and Code Inventory

Table 7: Complete resource inventory for reproducibility.

| **Resource** | **Path / URL** |
|---|---|
| Source code | `https://github.com/chemocalib/chemocalib` |
| Zenodo archive | `https://doi.org/10.5281/zenodo.21645890` |
| Docker image | `Dockerfile` in repository root |
| Conda environment | `environment.yml` |
| iJO1366 model | `https://bigg.ucsd.edu/models/iJO1366` |
| Multi-omics gen. | `chemocalib/data/loader.py` |
| 13C-MFA reference | `chemocalib/data/fluxome.py` |
| iJO1366 benchmark | `scripts/run_iJO1366_benchmark.py` |
| E-Flux2/SPOT script | `scripts/run_eflux2_spot_baselines.py` |
| Reliability diag. | `scripts/generate_reliability_diagram.py` |
| Figures | `scripts/generate_bioinformatics_figures.py` |
| Tutorial | `scripts/tutorial_5min.py` |
| Benchmark output | `output/ijo1366_benchmark.csv` |
| Notebooks | `notebooks/` (4 interactive tutorials) |
| Tests | `tests/` (8 test modules, 79 tests) |
